# Lysine demethylase 7a regulates the anterior-posterior development in mouse by modulating the transcription of *Hox* gene cluster

**DOI:** 10.1101/707125

**Authors:** Yoshiki Higashijima, Nao Nagai, Taro Kitazawa, Yumiko Kawamura, Akashi Taguchi, Natsuko Nakada, Masaomi Nangaku, Tetsushi Furukawa, Hiroyuki Aburatani, Hiroki Kurihara, Youichiro Wada, Yasuharu Kanki

## Abstract

Temporal and spatial colinear expression of the *Hox* genes determines the specification of positional identities during vertebrate development. Post-translational modifications of histones contribute to transcriptional regulation. Lysine demethylase 7A (Kdm7a) demethylates lysine 9 di-methylation of histone H3 (H3K9me2) and participates in the transcriptional activation of developmental genes. However, the role of Kdm7a during mouse embryonic development remains to be elucidated. Here, we show that *Kdm7a*^−/−^ mouse exhibits an anterior homeotic transformation of the axial skeleton, including an increased number of presacral elements. Importantly, posterior *Hox* genes (caudally from *Hox9*) are specifically downregulated in the *Kdm7a*^−/−^ embryo, which correlates with increased levels of H3K9me2. These observations suggest that Kdm7a controls the transcription of posterior *Hox* genes, likely via its demethylating activity, and thereby regulating the murine anterior-posterior development. Such epigenetic regulatory mechanisms may be harnessed for the proper control of coordinate body patterning in vertebrates.

## INTRODUCTION

*Hox* genes, which encode a family of homeodomain-containing transcription factors, are essential for the patterning of the anterior-to-posterior animal body axis during development. In mammals, the 39 *Hox* genes are divided into four clusters (*Hoxa*, *Hoxb*, *Hoxc*, and *Hoxd*) on four different chromosomes and consist of up to 13 paralogous groups. In each cluster, the *Hox* genes are arranged in tandem, from 3′ to 5′ (*Hox1* to *Hox13*). The 3′-paralogs are sequentially activated earlier than the 5′-paralogs along the anterior-posterior axis, a phenomenon that is called *Hox* temporal collinearity. This property of *Hox* expression confers special positional identities of the body segments, yet the underlying molecular mechanism is elusive. *Hox* transcription is switched on by retinoic acid signaling and morphogenic proteins, including Wnt and Fgf. Once the transcription starts, the newly activated *Hox* gene loci progressively cluster into a transcriptionally active chromatin compartment (Deschamps and Duboule, 2017; Deschamps and van Nes, 2005; Mallo and Alonso, 2013). Such transition in the spatial configuration coincides with the dynamics of chromatin histone marks, from a repressive state (tri-methylation of histone H3 lysine 27, H3K27me3) to a transcription-permissive state (tri-methylation of histone H3 lysine 4, H3K4me3) (Soshnikova and Duboule, 2009).

Polycomb group (PcG) proteins and the associated H3K27me3 mark maintain the state of transcriptional repression and gene silencing. Ezh2, a core component of the polycomb-repressive complex 2 (PRC2) is responsible for the methylation of H3K27me3. The *Hox* gene clusters are the best characterized PcG and H3K27me3 targets (Bracken et al., 2006; Pasini et al., 2010; Simon, 2010). Indeed, mutation of the PcG genes induces ectopic *Hox* expression, resulting in a posterior transformation of the axial skeleton in mouse (van Lohuizen, 1998). On the other hand, jumonji C (JmjC) domain-containing proteins, Utx and Jmjd3, specifically demethylate H3K27me2/3, and are involved in transcriptional activation of the *Hox* genes (Agger et al., 2007; Lan et al., 2007). Although catalytic action of Utx has been implicated in the regulation of expression of the *Hox* genes during zebrafish development (Lan et al., 2007), it has been recently demonstrated that mouse with catalytically-inactive Jmjd3, but not an Utx mutant, exhibits anterior homeotic transformation associated with a downregulation of *Hox* genes (Naruse et al., 2017).

Di-methylation of histone H3 lysine 9 (H3K9me2), another repressive histone mark, is methylated by SET domain-containing proteins, G9a (encoded by *Ehmt2*) and GLP (encoded by *Ehmt1*) (Ogawa et al., 2002; Tachibana et al., 2001; Tachibana et al., 2002). H3K9me2 is the most abundant heterochromatic histone modification, and covers large genomic domains in differentiated cells and in embryonic stem (ES) cells (Lienert et al., 2011; Poleshko et al., 2017; Wen et al., 2009). These domains are specifically associated with lamina-associated domains (LADs), characterized as transcriptionally-repressive heterochromatin located within the nuclear peripheral region. A negative correlation between H3K9me2 deposition and gene expression is observed therein. During mouse embryogenesis, repressed *Hox* genes labeled by the H3K27me3 marks are located at a spatial domain distinct from the peripheral LADs (Vieux-Rochas et al., 2015). Consistently, the association of genomic occupancies of H3K9me2 and H3K27me3 is mutually exclusive during differentiation of the mouse ES cells (Lienert et al., 2011; Zylicz et al., 2015). Indeed, to the best of our knowledge, the functional relationship between *Hox* gene expression and the H3K9me2 histone mark has not been ruled out to date.

Another JmjC domain-containing protein, lysine demethylase 7A (Kdm7a), also known as Jhdm1d, contains a plant homeo domain (PHD), and is responsible for the demethylation of H3K9me2 and H3K27me2 (Huang et al., 2010b; Tsukada et al., 2010). Kdm7a is predominantly expressed in mouse brain tissues. Inhibition of a Kdm7a ortholog in zebrafish leads to developmental brain defects. In mammalian neuronal cells, Kdm7a binds to the gene locus of *follistatin*, an antagonist of activin, which plays an important role in brain development. Kdm7a depletion suppresses the transcription of the gene, in association with increased levels of demethylated H3K9 and H3K27 (Tsukada et al., 2010). In addition, Kdm7a promotes neural differentiation of mouse ES cells by transcriptional activation of Fgf4, a signal molecule implicated in neural differentiation. Knockdown of *Kdm7a* decreases Fgf4 expression, which correlates with the enriched coverage of both H3K9me2 and H3K27me2 (Huang et al., 2010b). Furthermore, Kdm7a ortholog is predominantly expressed in epiblast cells of the primitive streak and promotes neural induction in an early chick embryo (Huang et al., 2010a). However, the biological role of Kdm7a during mouse development has not yet been reported.

Here, we report that *Kdm7a*^−/−^ mutant mouse exhibits anterior homeotic transformation of the axial skeleton and downregulation of the transcription of posterior *Hox* genes during embryogenesis. Importantly, these changes in gene expression are associated with increased H3K9me2 levels at the relevant posterior *Hox* loci. These observations demonstrate an essential role of Kdm7a on chromatin structure and *Hox* gene regulation *in vivo*. Further, they provide evidence for the role of epigenetic histone mark H3K9me2 in the maintenance of *Hox* gene regulation during embryonic development in mouse.

## RESULTS

### Construction of a *Kdm7a*^−/−^ mouse by CRISPR/Cas9-mediated gene targeting

To disrupt the enzyme function of Kdm7a, we used a CRISPR/Cas9-based strategy to introduce a frameshift mutation at the start of the JmjC domain in *Kdm7a*. Because there are no suitable protospacer-adjacent motif (PAM) sequences in exon5 of the region encoding the JmjC domain, we designed single guide RNAs (sgRNAs) located in exon6 of the region encoding the JmjC domain (Figure 1A). To determine the optimal sgRNA sequence, we co-transfected HeLa cells with the pCAG-EGxxFP-target and pX330-sgRNA plasmids. We monitored the reconstituted enhanced green fluorescent protein (EGFP) fluorescence 48 h after transfection. Cetn1 was used as a positive control (Mashiko et al., 2013). Although both sgRNA867 and sgRNA868 effectively cleaved the target site of pCAG-EGxxFP-Kdm7a, sgRNA868 worked slightly better than sgRNA867. We therefore selected sgRNA868 for further *in vivo* genome editing (Figure 1B).

**Figure 1.**
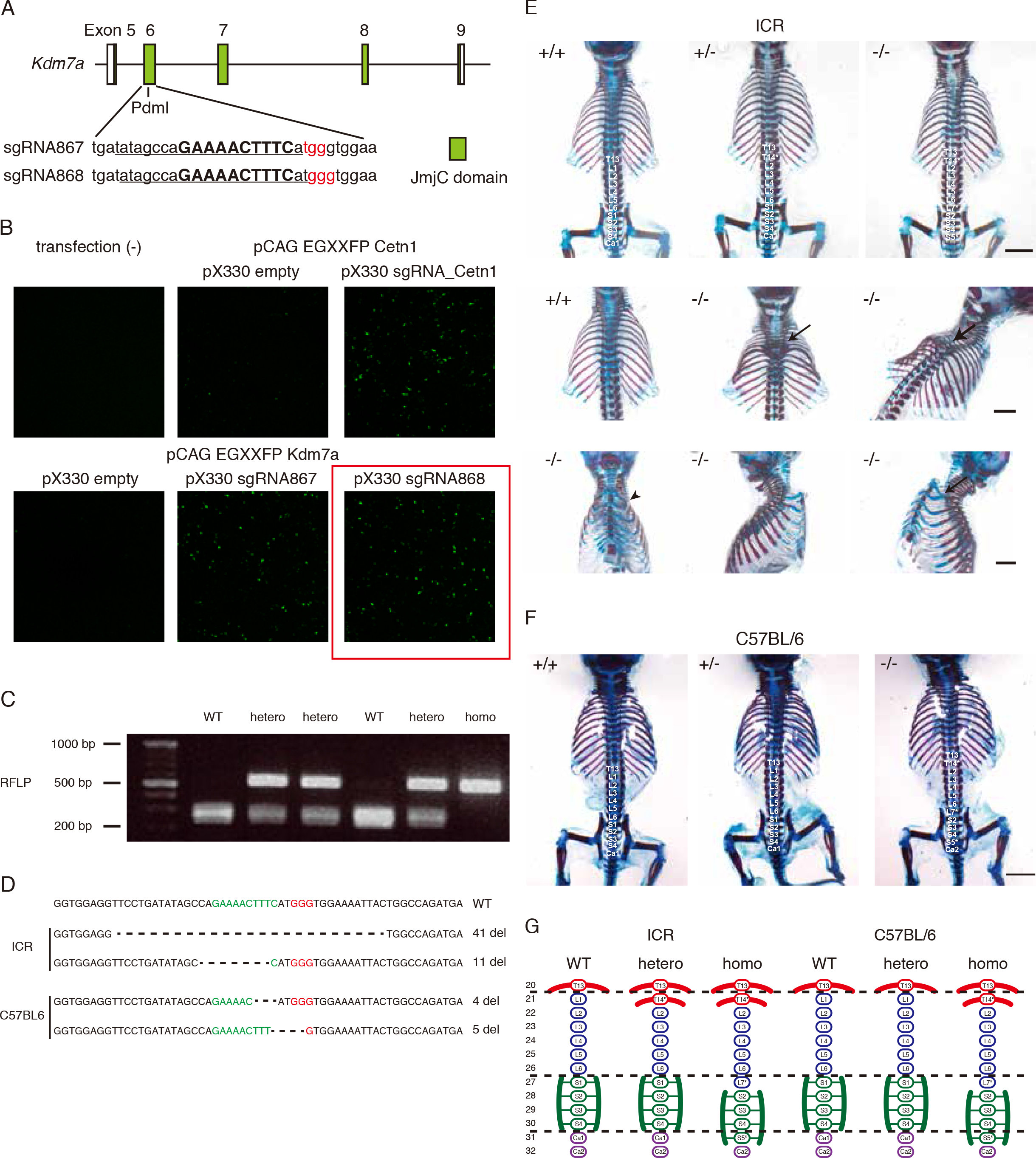
Kdm7a regulates the anterior-posterior patterning of the axial skeleton in mouse. (A) Schematic of the Cas9/sgRNA-targeting sites in *Kdm7a*. The sgRNA-targeting sequence is underlined, and the protospacer-adjacent motif (PAM) sequence is labeled in red. The restriction sites in the target regions are bold and capitalized. Restriction enzymes used for restriction fragment length polymorphism (RFLP) are shown, and JmjC domain is shown as green boxes. (B) Validation of double-strand break (DSB)-mediated homology-dependent repair by reconstitution of the enhanced green fluorescent protein (EGFP). The approximately 500-bp genomic fragment containing the sgRNA target sequence (*Cetn1* for positive control or *Kdm7a* for sgRNA validation) was inserted in pCAG-EGxxFP target plasmid (Mashiko et al., 2013). The pX330 plasmid contains a humanized Cas9 expression cassette and an sgRNA expression cassette. The sgRNA targeting *Cetn1* or *Kdm7a* (Cetn1: sgRNA_Cetn1, KDM7A: sgRNA867 or sgRNA868) was cloned into the pX330 plasmid. Both pCAG-EGxxFP and pX330 plasmid were co-transfected into the HeLa cells. When the target sequence was digested by sgRNA-guided Cas9 endonuclease, homology dependent repair (HR, homologous recombination; SSA, single-strand annealing) resulted in the reconstitution of the EGFP expression cassette. (C) RFLP analysis. *Kdm7a* PCR products were digested with PdmI. Representative RFLP result is shown. (D) The sequence of mutant alleles in *Kdm7a* KO mice from ICR or C57BL/6 backgrounds. PAM sequence and the restriction site are labeled in red and green, respectively. (E) Patterning defects in the axial skeleton of *Kdm7a* KO ICR background mouse. In *Kdm7a*^−/−^ mice, the 1^st^ lumbar (L1), the 1^st^ sacral (S1) and the 1^st^ coccygeal (Co1) vertebrae were transformed into thoracic (T14*), lumbar (L7*) and sacral (S5*) elements, respectively. In *Kdm7a*^+/−^ mice, only L1 was transformed into thoracic (T14*) element. * indicates transformed elements. Dorsal views of the vertebrosternal ribs in the wild-type (center left) and *Kdm7a*^−/−^ (center middle and right) mice are shown. The arrows indicate the presence of cervical vertebral phenotypes in thoracic vertebrae. Ventral (lower left), left lateral (lower middle), and right lateral (lower right) views of the vertebrosternal ribs in *Kdm7a*^−/−^ mice are shown. The arrowhead indicates the fusion of ribs. (F) Homeotic transformation in the axial skeleton of *Kdm7a* KO C57BL/6 mouse. In *Kdm7a*^−/−^ mice, L1, S1 and Co1 were transformed into thoracic (T14*), lumbar (L7*) and sacral (S5*) elements, respectively. *Kdm7a*^+/−^ background showed no patterning defects. * indicates transformed elements. (G) Summary of the patterning defects identified across *Kdm7a* mutant alleles in the ICR and C57BL/6 backgrounds. An asterisk indicates a homeotic transformation of the vertebral element.

Accordingly, we co-injected *Cas9* mRNA with sgRNA868 into pronuclear stage one-cell mouse embryos. The blastocysts derived from the injected embryos were then transplanted into foster mothers and newborn pups were obtained. Mice carrying the targeted mutations (chimera mice) were crossbred with the wild type and heterozygous mice were obtained. Representative results of the restriction fragment length polymorphism (RFLP) analysis are shown in Figure 1C. We then amplified the Kdm7a-targeted regions by polymerase chain reaction (PCR), and subcloned and sequenced the PCR products, to confirm that the tested mice carried the mutant alleles with small deletions at the target site (Figure 1D). Because the phenotypes of mutant mice may differ between genetic backgrounds (Doetschman, 2009), we generated *Kdm7a* mutant mice in both, ICR and C57BL/6 backgrounds. These mutant mice were used for subsequent analysis *in vivo*.

### Kdm7a regulates the anterior-posterior patterning of the axial skeleton in mouse

The *Kdm7a* mutant newborns appeared grossly normal. Considering that epigenetic factors, including histone demethylases, are associated with the anterior-posterior patterning (Hong et al., 2018; Jansz et al., 2018; Naruse et al., 2017; Terranova et al., 2006), we investigated whether Kdm7a plays a role in the animal body patterning. To this end, we generated whole-mount skeletal preparations of postnatal day 1 wild-type and *Kdm7a* mutant mice. As anticipated, all wild-type mice demonstrated the normal configuration of the axial skeleton, with 7 cervical, 13 thoracic, 6 lumbar and 4 sacral vertebrae (five out of five animals and 12 out of 12 animals from the ICR and C57BL/6 background, respectively) (Figure 1E-G, Tables 1 and 2). By contrast, in all *Kdm7a*^−/−^ mice displayed an anterior homeotic transformation of vertebral elements. The 1^st^ lumber vertebra (L1) transformed into the thoracic element (T14*) gaining ectopic ribs, and the 1^st^ sacral (S1) and coccygeal (Co1) vertebrae showed transformation to lumbar (L7*) and sacral (S5*) identities with the loss and gain of connections to the pelvic girdle, respectively (11 out of 11 animals, and eight out of eight animals from the ICR and C57BL/6 background, respectively) (Figure 1E-G, Tables 1 and 2). Of note, even heterozygous mutant mice from ICR background exhibited an anteriorization of L1 into the thoracic element (T14*) (four out of five animals), while those from C57BL/6 background showed no difference from the normal vertebral disposition (eleven out of eleven animals) (Figure 1E-G, Tables 1 and 2). In addition, *Kdm7a*^−/−^ mice from the ICR but not C57BL/6 background displayed varying changes in vertebral morphology in the vicinity of the thoracic vertebra and ribs (i.e., the presence of cervical vertebral phenotypes in the thoracic vertebrae and the fusion of ribs) (four out of eleven animals) (Figure 1E), suggesting that the ICR genetic background has a much stronger influence on the anterior-posterior patterning in *Kdm7a* mutant mouse than the C57BL/6 background. Nonetheless, *Kdm7a* mutant mice showed anterior homeotic transformation of the axial skeleton regardless of the genetic background (schematized in Figure 1G). Considering that an ICR mouse is a non-inbred strain and genetically heterogeneous, we decided to conduct further detailed genetic analysis involving C57BL/6 mice.

**Table 1:** Axial skeletal phenotypes of *Kdm7a* mutant mice (ICR background).

|  | Wild-type | Hetero | Homo |
| --- | --- | --- | --- |
| Animal number | 5 | 5 | 11 |
| **Vertebral pattern** |  |  |  |
| T:L:S = 13:6:4<br>(T1-T13, L1-L6, S1-S4) | 100% | 0% | 0% |
| T:L:S = 14:5:4<br>(T1-T14*, L2-L6, S1-S4) | 0% | 80% | 0% |
| T:L:S = 14:6:4<br>(T1-T14*, L2-L7*, S2-S5*) | 0% | 20% | 100% |

**Table 2:** Axial skeletal phenotypes of *Kdm7a* mutant mice (C57BL/6 background).

|  | Wild-type | Hetero | Homo |
| --- | --- | --- | --- |
| Animal number | 12 | 11 | 8 |
| **Vertebral pattern** |  |  |  |
| T:L:S = 13:6:4<br>(T1-T13, L1-L6, S1-S4) | 100% | 100% | 0% |
| T:L:S = 14:5:4<br>(T1-T14*, L2-L6, S1-S4) | 0% | 0% | 0% |
| T:L:S = 14:6:4<br>(T1-T14*, L2-L7*, S2-S5*) | 0% | 0% | 100% |

### Kdm7a is involved in the regulation of *Hox* gene expression

Loss-of-function of the murine *Hox* genes classically causes an anterior homeotic transformation (Wellik and Capecchi, 2003). Hence, we next examined the expression of *Hox* genes during embryogenesis by RNA sequencing (RNA-Seq). In the experiments, wild-type and *Kdm7a* mutant embryos at E9.5 and E10.5 were divided at the level of the otic vesicle (hereafter referred to as the “trunk”) (beige-colored in Figure 2A). The embryonic trunk at this developmental stage is a region in which the *Hox* genes are predominantly expressed (Deschamps and Duboule, 2017). Data for three biological replicates of RNA-Seq analyses were highly correlated (data not shown). The analysis revealed that 73 and 2 genes were differentially expressed between the wild-type and *Kdm7a*^−/−^ embryos at E9.5 and E10.5, respectively (padj<0.05; Figure 2B and C, Supplementary Table 1). A decreased expression of *Kdm7a* was detected in *Kdm7a*^−/−^ embryo, which was probably associated with a nonsense-mediated mRNA decay (Popp and Maquat, 2016). Of note, many of the genes, including *Hox*, were downregulated in the *Kdm7a*^−/−^ embryo, suggesting the possible role of Kdm7a in transcriptional activation (Figure 2B and C). This was consistent with a previous study showing that genetic ablation of H3K9me2 methyltransferase G9a resulted in the activation of many genes (upregulation of 147 and downregulation of 33 transcripts) (Zylicz et al., 2015). As anticipated, gene ontology (GO) analysis and Ingenuity Pathway Analysis (IPA) of 73 differentially expressed genes, excluding *Kdm7a*, revealed a significant enrichment of the “skeletal system development”, “anterior/posterior pattern specification”, and “development of body axis” processes (Figure 2D and E). In addition, the characteristics of these differentially expressed genes were related to the component “nucleus”, the function “sequence-specific DNA binding”, and the sequence domain “HOX” and “Homeobox, conserved site”. This indicated that Kdm7a participates in the regulation of the developmental transcription factors, including Hox (Figure 2D-F). Interestingly, when we focused on all (39) *Hox* genes, we observed that the posterior *Hox* genes were downregulated, while there were no differences in the expression in the anterior *Hox* genes in the *Kdm7a*^−/−^ embryo compared with the wild type (Figure 2G). Quantitative PCR (qPCR) analysis confirmed that the expression of the majority of posterior *Hox* genes (*Hoxb6*; *c9*; *d9*; *a10*; *c10*; *d10*; *a11*; *c11*; *d11*; *d12*; and *d13* for E9.5; and *Hoxd8*; *c10*; *a11*; *c11*; *d12*; *a13*; *c13*; and *d13* for E10.5) was significantly decreased in the *Kdm7a*^−/−^ embryo, and this was evident in an E9.5 comparison with E10.5 (Figure 2H). Collectively, these findings support a functional role of Kdm7a-mediated transcriptional control, especially of the posterior *Hox* genes.

**Figure 2.**
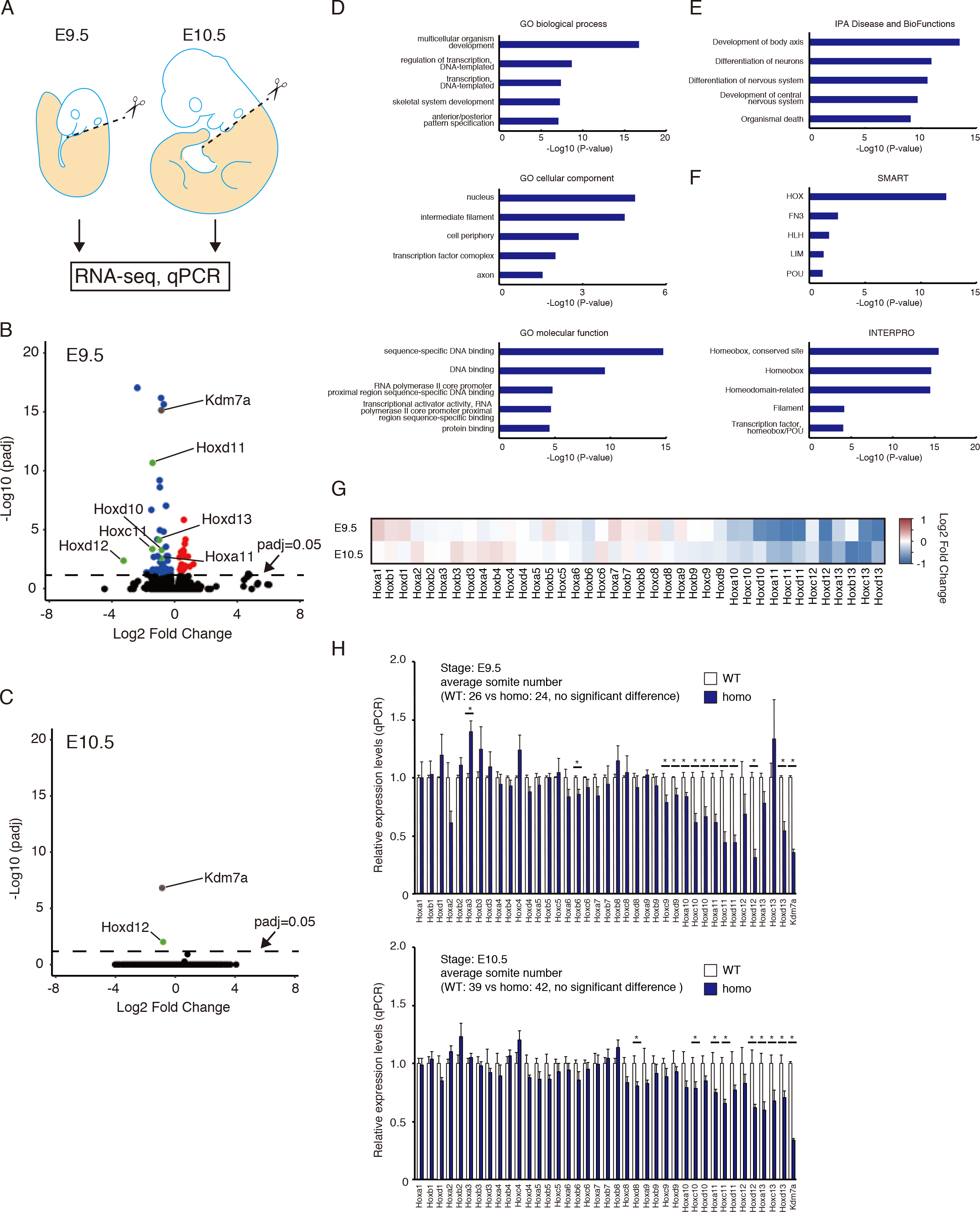
Kdm7a is involved in the regulation of *Hox* gene expression. (A) Schematic of E9.5 and E10.5 mouse embryo microdissection. The posterior part of the embryo (beige; referred to as the “trunk”) was used for RNA sequencing (RNA-Seq) and quantitative polymerase chain reaction (qPCR) analysis. (B and C) Volcano plots showing differentially expressed genes in the wild-type and *Kdm7a*^−/−^ embryos (*n*=3 for each genotype) at E9.5 (B) and E10.5 (C). The X- and Y-axes indicate the log2 fold-change and −log10 adjusted *P*-value (padj) produced by DESeq2, respectively. Genes with padj≤0.05 are indicated as red (increase) and blue (decrease) spots. (D-F) Gene ontology (GO) analysis (D), Ingenuity Pathway Analysis (IPA) (E), and domain prediction analysis (InterPro and SMRT) (F) of 73 differentially expressed genes in *Kdm7a*^−/−^ mouse, as determined in Figure 2B. The *P*-values for each category are shown in the bar graphs. (G) Heat maps showing the log2 fold-change expression differences (determined by DESeq2) in *Hox* genes between the wild-type and *Kdm7a*^−/−^ embryos at E9.5 and E10.5. Red to blue coloring indicates the fold change. (H) qPCR analysis comparing the expression of *Hox* genes between wild-type and *Kdm7a*^−/−^ embryos at E9.5 (top; *n*=5 for each genotype) and E10.5 (bottom; *n*=4 for each genotype). The average number of somites in the wild-type and *Kdm7a*^−/−^ was 26 and 24 at E9.5, respectively, and 39 and 42 at E10.5, respectively (there were no statistically significant differences between the wild-type and *Kdm7a*^−/−^ embryos). Data are shown as means ± SE. \**P*<0.05 compared with the wild type.

### H3K9me2 methylation is involved in the regulation of *Hox* genes

Kdm7a-mediated demethylation of the repressive histone mark H3K9me2 correlates with active gene expression (Huang et al., 2010b; Tsukada et al., 2010). Therefore, we hypothesized that the transcriptional activation/repression of *Hox* genes during the anterior-posterior patterning would be associated with decreased/increased levels of H3K9me2. Because the posterior *Hox* genes are not activated in the developmental brain (Keynes and Krumlauf, 1994), we first compared the epigenetic landscape of H3K9me2 in the developmental brain (hereafter referred to as the “head”) with the trunk (Figure 3A). Although the expression of *Hox* genes is more subject to change at E9.5 than E10.5, we selected the latter time point because of the requirement for large numbers of cells for the chromatin immunoprecipitation (ChIP) analysis. As expected, ChIP-Seq analysis revealed increased levels of H3K9me2 at almost all *Hox* genes in the head regions compared with the trunk (Figure 3B). At a representative *Hoxa* cluster locus, ChIP-Seq signals for H3K9me2 were broadly increased in the embryonic head, which coincided with the decreased signals for the active histone mark H3K4me3 (Figure 3C). ChIP followed by qPCR revealed an increase in H3K9me2 levels and a decrease in H3K4me3 levels in the vicinity of the transcription start site (TSS) of *Hoxa3* and *Hoxa13*, but no changes at *actin beta* (*Actb*) (Figure 3D). We next examined whether ablation of Kdm7a affected H3K9me2 levels at the *Hox* genes in the trunk regions (Figure 3E). In accordance with the mRNA expression data (Figure 2G and H), H3K9me2 coverage was enriched at the posterior *Hox* genes in the *Kdm7a*^−/−^ embryo in comparison with the wild type (Figure 3F). We observed high occupancy of H3K9me2 at the *Hoxa* locus in the *Kdm7a*^−/−^ embryonic trunk, but no differences in the levels of H3K4me3 (Figure 3G). Consistently, ChIP followed by qPCR showed an increase in H3K9me2 levels in the vicinity of the TSS of *Hoxa3* and *Hoxa13*, but no changes at *Actb* (Figure 3H). Taken together, these observations suggest the possibility that Kdm7a-mediated regulation of the repressive histone mark H3K9me2 might be involved in transcriptional activation of the *Hox* genes.

**Figure 3.**
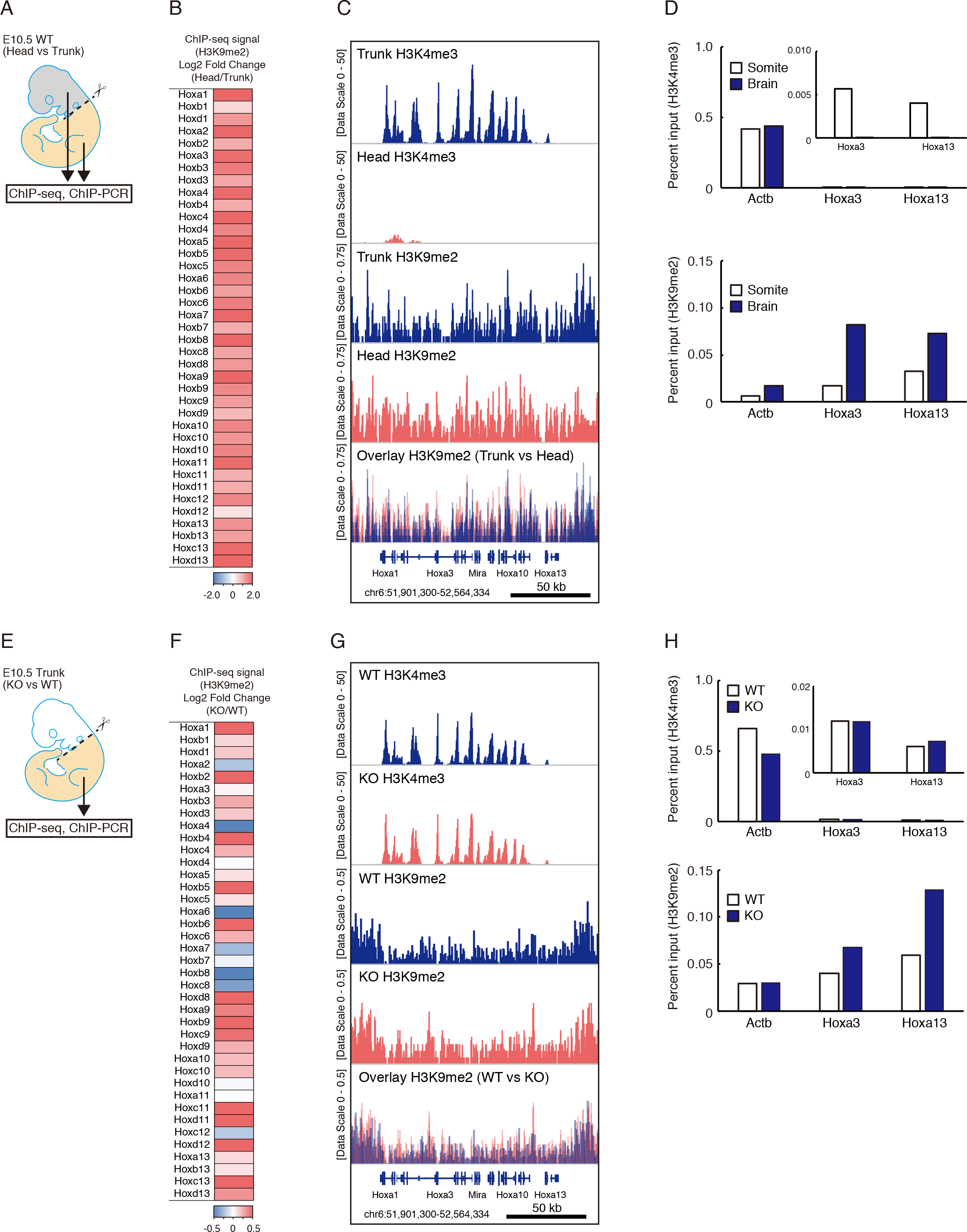
H3K9me2 methylation is involved in the regulation of *Hox* genes. (A) The developmental brain (referred to as the “head”) and trunk from the wild-type embryo at E10.5 (colored grey and beige, respectively) were used for chromatin immunoprecipitation (ChIP)-Seq and ChIP-qPCR. (B) Heat maps showing the log2 fold-change in H3K9me2 ChIP-Seq signals in the *Hox* genes in the head and trunk regions of the wild-type embryo. Red to blue coloring indicates the fold change. (C) Gene tracks of ChIP-Seq signals for H3K4me3 and H3K9me2 close to the *Hoxa* cluster in the head and trunk of the wild-type embryo. ChIP-Seq signals were visualized by using Integrative Genomics Viewer (http://software.broadinstitute.org/software/igv/). (D) ChIP-qPCR of H3K4me3 (top) and H3K9me2 (bottom) at the *Actb*, *Hoxa3*, and *Hoxa13* transcription start site (TSS) in the head and trunk regions from the wild-type embryo, normalized to input. Data are representative of two independent experiments. (E) Developmental trunks from the wild-type or *Kdm7a*^−/−^ embryos at E10.5 (beige) were used for ChIP-Seq and ChIP-qPCR. (F) Heat maps showing the log2 fold-change in H3K9me2 ChIP-Seq signals in the *Hox* genes between the trunk regions from the wild-type and *Kdm7a*^−/−^ embryos. Red to blue coloring indicates the fold change. (G) Gene tracks of ChIP-Seq signals for H3K4me3 and H3K9me2 close to the *Hoxa* cluster in the trunk-region of the wild-type and *Kdm7a*^−/−^ embryos. ChIP-Seq signals were visualized by Integrative Genomics Viewer. (H) ChIP-qPCR of H3K4me3 (top) and H3K9me2 (bottom) at the *Actb*, *Hoxa3*, and *Hoxa13* TSS in the trunk regions of the wild-type and *Kdm7a*^−/−^ embryos, normalized to input. Data are representative of two independent experiments.

## DISCUSSION

In the last decades, histone-modifying enzymes have become recognized as the key players of the development, differentiation, and various diseases. Kdm7a, a histone demethylase for H3K9me2 and H3K27me2, is reportedly involved in neural differentiation of mouse ES cells and in brain development in zebrafish (Huang et al., 2010b; Tsukada et al., 2010). In addition, Kdm7a is highly induced in cancer cells in response to nutrient starvation and is associated with tumor suppression, by modulating tumor angiogenesis (Osawa et al., 2011). However, its role in mouse development has not been elucidated. We report here the generation and characterization of a previously undescribed *Kdm7a* mouse mutant. We provide the evidence that Kdm7a is involved in the activation of the posterior *Hox* gene expression, and subsequent patterning of the anterior-posterior body axis, *in vivo*. Since we observed increased levels of H3K9me2 at the relevant posterior *Hox* loci in the *Kdm7a* mutant embryo, we propose that Kdm7a modulates the developmental *Hox* gene activation by regulating the repressive histone mark H3K9me2.

Many studies support the notion that the mammalian *Hox* genes are targets of PcG proteins and their associated H3K27me3, indicating the essential role of H3K27me3 in the silencing of *Hox* gene expression (Lonfat et al., 2014; Narendra et al., 2015; Soshnikova and Duboule, 2009; Vieux-Rochas et al., 2015). In the mouse ES cells, H3K27me3 covers the entire *Hox* clusters, in which the *Hox* genes are transcriptionally repressed. Further, during collinear activation of the *Hox* genes, H3K27me3 marks at the *Hox* cluster loci are progressively diminished in the sequence of transcriptional activation of the *Hox* genes (Soshnikova and Duboule, 2009). Accordingly, posterior transformation and an increased expression of the *Hox* genes are commonly seen in mice lacking PcG proteins (van Lohuizen, 1998). For example, a mouse with mutation in the *Mel18* gene, also known as the polycomb group ring finger 2 gene, exhibits a posteriorizing shift of body axis (e.g., the loss of rib in the thoracic vertebrae and ectopic ribs in the cervical vertebrae) (Akasaka et al., 1996; Suzuki et al., 2002). Furthermore, *Jmjd3* mutant mouse, in which the protein’s H3K27 demethylation domain is disrupted, exhibits an anterior homeotic transformation (e.g., the gain of rib in the lumbar vertebra), which is associated with the downregulation of *Hox* genes (Naruse et al., 2017). Hence, although we observed the involvement of H3K9me2 in the transcriptional regulation of *Hox* genes, we could not rule out the possible association of H3K9me2 with the H3K27me3-mediated repression mechanisms.

While both H3K9me2 and H3K27me3 are involved in facultative heterochromatinization during the development, a limited overlap between H3K9me2 and H3K27me3 targets has been suggested (Trojer and Reinberg, 2007). Indeed, during the early postimplantation development, only a few genes are differentially expressed between the *Ehmt2*^−/−^ and *Ezh2*^−/−^ mutant embryos, which is in line with H3K9me2 and H3K27me3 being linked to distinct repressive chromatin states (Zylicz et al., 2015). Accordantly, genome-wide analysis of the differentiation of mouse ES cells revealed that the occurrence of H3K9me2 and H3K27me3 is mutually exclusive, with relatively sharp boundaries between the two marks (Lienert et al., 2011). Another example concerns “germline genes”, which are crucial for the progression of the primordial germ cell to meiosis in the female and for transposon repression in the male. These genes are silenced by both H3K9me2 and H3K27me3 in the mouse ES cells, and are progressively activated in association with decreased H3K9me2 marks, along with the specification and development of the primordial germ cell (Kurimoto et al., 2015). In addition, deposition of both H3K9me2 and H3K27me3 in the vicinity of the genomic region of *Pax5*, regulated by PcG proteins, is simultaneously decreased by *Ehmt2* knockout in the mouse ES cells (Wen et al., 2009). Furthermore, siRNA knockdown of *Kdm7a* in neuronal cells led to increased levels of not only H3K9me2 but also H3K27me3 at the *follistatin* locus (Tsukada et al., 2010). Taken together, H3K9me2 and H3K27me3 normally have independent functions, as observed in mouse ES cells. Nevertheless, cooperative transcriptional control by H3K9me2 and H3K27me3 could occur under certain conditions.

In mouse ES cells, as well as in the early embryo, H3K27me3-marked *Hox* clusters are located at a relatively central and active nuclear environment, away from the peripheral LADs, in which H3K9me2 marks are enriched (Vieux-Rochas et al., 2015). Despite this, we here demonstrated a potential mechanism by which H3K9me2 controls the *Hox* gene expression during mouse development. The following model might be proposed to explain the discrepancy of the localization of the H3K9me2 and H3K27me3 marks. Several groups have reported that abrogation of H3K9me2 by *G9a* knockdown leads to a loss of peripheral interactions of its associated genes (Bian et al., 2013; Chen et al., 2014; Harr et al., 2015). Furthermore, it has been recently shown that mechanical strain causes defective chromatin anchoring at the nuclear lamina and a switch from H3K9me2 to H3K27me3 occupancy at their associated genomic loci in the epidermal stem cells (Le et al., 2016). Further, H3K27me3 coverage in *Hox*-negative differentiated cells is relatively enriched compared to that in mouse ES cells where the *Hox* genes are also not transcribed (Deschamps and Duboule, 2017; Noordermeer et al., 2011). Hence, Kdm7a-mediated demethylation of H3K9me2 might be possibly involved in the transition of the *Hox* clusters to the H3K27me3-enriched, more active nuclear environment and a subsequent transcriptional control of the *Hox* genes, although further studies are needed to investigate the mechanism by which H3K9me2 and H3K27me3 cooperatively mediate regulation of the *Hox* genes.

In the present study, we showed that the posterior *Hox* genes are preferentially downregulated in *Kdm7a* mutant embryos with an occurrence of, accordingly, anterior transformation (i.e., ectopic ribs; T13 to T14). Jansz et al. (2018) have recently reported that the structural maintenance of chromosomes (SMC) hinge domain containing 1(Smchd1), a noncanonical SMC protein, is involved in the regulation of the chromosomal conformation interactions of the *Hox* cluster and *Hox* gene silencing. Genetic ablation of Smchd1 results in an activation of the posterior *Hox* genes without affecting the anterior *Hox* genes at the same developmental stage as that of embryos analyzed in the current study, and a subsequent posterior transformation (i.e., loss of ribs; T13 to T12), representing an inverse phenotype to that of the *Kdm7a* mutant (Jansz et al., 2018). More recently, we reported that KDM7A-mediated demethylation of H3K9me2 is critical for the early inflammatory responses and is associated with conformational changes of the chromatin between inflammation-induced enhancers in human endothelial cells (Higashijima et al., 2018). Collectively, Kdm7a might likely play a role in chromosomal conformation for posterior *Hox* gene regulation at a certain time of development.

Recent findings have suggested a non-catalytic function of histone-modifying enzymes, especially in tumorigenesis. UTX-mediated chromatin remodeling suppresses acute myeloid leukemia via a noncatalytic inverse regulation of the oncogenic and tumor-suppressive transcription factor programs (Gozdecka et al., 2018). In addition, a non-enzymatic function of SETD1A, a methyltransferase of H3K4, regulates the expression of genes involved in DNA damage response and is required for the survival of acute myeloid leukemia cells (Hoshii et al., 2018). In the present study, we showed that H3K9me2 occupancy is enriched in the *Ho*x-negative developmental brain and increased in the posterior part of the *Kdm7a* mutant embryo. Hence, we believe that the catalytic activity of Kdm7a possibly plays an important role in the transcriptional control of *Hox* genes during embryogenesis. Nonetheless, experiments involving a catalytically inactive mutant will be required to clarify this point.

In conclusion, the presented data establish an important *in vivo* role of Kdm7a in the anterior-posterior axial development. Kdm7a regulates the transcription of *Hox* genes most likely by the demethylation of the repressive histone mark H3K9me2. Such systems might be essential for the proper control of coordinate body patterning in vertebrate development. Currently, studies focusing on the role of H3K9me2 during embryogenesis are limited (Au Yeung et al., 2019), and further studies are warranted to understand the mechanisms through which H3K9me2 mediates transcriptional regulation of the developmental genes, including *Hox*.

## ACKNOWLEDGEMENTS

We thank Shiro Fukuda and Shogo Yamamoto (The University of Tokyo) for bioinformatics analysis. pX330 plasmid vectors was kindly gifted from Dr. Masahito Ikawa (Osaka University). This work was supported by a Grant-in-Aid for JSPS Postdoctoral Fellows (to Y.H.); a Grant-in-Aid for Young Scientists (B) 17K15991 (to Y.H.); a Grant-in-Aid for Young Scientists (A) 26710013 (to Ya.K.); a Grant-in-Aid for Scientific Research on Innovative Areas (Research in a Proposed Research Area) 25125707 (to Ya.K.); a Grant-in-Aid for Challenging Exploratory Research [26670397 (to Ya.K.) and 16K15438 (to Ya.K.)]; a Fund for the Promotion of Joint International Research (Fostering Joint International Research) 15KK0251 (to Ya.K.); a Research Grant from Nanken-Kyoten, TMDU (to Y.H., Ya.K., Y.W., and T.F.); a Research Grant from Takeda Science Foundation (to Ya.K.); a Research Grant from the Japan Heart Foundation (to Ya.K.); a Research Grant from MSD Life Science Foundation (to Ya.K.); a Research Grant from Uehara Memorial Foundation (to Ya.K.); a Research Grant from SENSHIN Medical Research Foundation (to Ya.K.); and a Research Grant from Kowa Life science Foundation (to Ya.K.).

## AUTHOR CONTRIBUTIONS

Y.H. T.K. Yu.K. and Ya.K designed the research strategies; Y,H. T.K. Yu.K. A.T. Nat.N, and Ya.K. performed the experiments; Y,H. Nao.N. and Ya,K performed the bioinformatic analyses; Y,H. Nao.N. T.K. Yu.K. M.N. H.K. H.A. Y.W. T.F. and Ya.K analyzed the data; and Y,H. Nao.N. T.K. and Ya.K. wrote the manuscript.

## METHOD DETAILS

### Mice

All mouse experiments were approved by The University of Tokyo Animal Care and Use Committee (approval number H28-1). The animals were housed in individual cages in a temperature- and light-controlled environment, and had *ad libitum* access to chow and water.

### Cell lines

Human cervical cancer cell line, HeLa, was purchased from ATCC (Manassas, VA) and grown and passaged every 2 or 3 days in DMEM (nacalai tesque, Kyoto, Japan), supplemented with 1% penicillin/streptomycin (Wako, Osaka, Japan) and 10% FBS (Thermo Fisher Scientific, Waltham, MA). The cells were cultured at 37 °C and in a 5% CO_2_ atmosphere in a humidified incubator.

### Plasmids and mRNA preparation

The pCAG-EGxxFP (Mashiko et al., 2013) plasmid was a kind gift from Dr. M Ikawa (The University of Osaka). Genomic fragments (approximately 500-bp) containing the sgRNA target sequence were PCR-amplified and placed between the EGFP-encoding fragments. Plasmids expressing both hCas9 and sgRNA were prepared by inserting synthetic oligonucleotides (Hokkaido System Science, Hokkaido, Japan) at the BbsI site of pX330 (http://www.addgene.org/42230/) (Cong et al., 2013). Plasmids pCAG-EGxxFP, harboring the sgRNA target sequence of *Cetn1*, and pX330, containing sgRNA targeting *Cetn1*, were also kindly gifted from Dr. M Ikawa (Mashiko et al., 2013). The p3s-Cas9HC plasmid (https://www.addgene.org/43945/) was used to generate hCas9 mRNA. The plasmid for producing sgRNA was prepared by inserting synthetic oligonucleotides (Hokkaido System Science) at the BsaI site of DR274 (https://www.addgene.org/42250/). RNA was synthesized from the XbaI-digested p3s-Cas9HC plasmid by using mMESSAGE mMACHINE T7 ULTRA transcription kit (Thermo Fisher Scientific) in accordance with manufacturer’s protocol. RNA was synthesized from the DraI-digested DR274 plasmid by using MEGAshortscript™ T7 transcription kit (Thermo Fisher Scientific) in accordance with manufacturer’s protocol. The hCas9 mRNA and sgRNAs were purified by phenol chloroform-isoamyl alcohol extraction and isopropanol precipitation, followed by spin column chromatography using NANOSEP MF 0.2 μm (Thermo Fisher Scientific). The PCR primers and oligonucleotide sequences for the constructs are listed in Supplementary Table 2.

### Transfection procedure

For the experiment, 250 ng of pCAG-EGxxFP-target was mixed with 250 ng of pX330 harboring the sgRNA sequences, and the mixture was used to transfect 1 × 10^5^ HeLa cells in a well of a 24-well plate using the Lipofectamine® LTX reagent (Thermo Fisher Scientific), according to the manufacturer’s protocol. The EGFP fluorescence was observed under a confocal microscope (C2^+^ Confocal Microscope System; Nikon, Tokyo, Japan) 48 h after the transfection.

### Pronuclear injection

ICR and C57BL/6 female mice were superovulated and mated with ICR and C57BL/6 males, respectively, and fertilized eggs were collected from the oviduct. Then, the hCas9 mRNA (0.05 μg/μl) and sgRNAs (0.05 μg/μl) were co-injected into pronuclear-stage eggs. The eggs were cultivated in kSOM overnight and then transferred into the oviducts of pseudopregnant ICR females.

### Genotyping

Mouse genomic DNA samples were prepared from tail biopsies. PCR was performed using *Kdm7a*-specific primers to amplify the sgRNA target site (Supplementary Table 2), and under the following cycling conditions: 95°C for 10 min; followed by 40 cycles of 95°C for 20 s, 60°C for 20 s, and 72°C for 30 s; incubation step at 72°C for 7 min; and hold at 4°C. BMS BIOTAQ™ DNA polymerase (Nippon Genetics Co. Ltd, Tokyo, Japan) was used for PCR reactions. The *Kdm7a* PCR product was digested with XmnI (New England Biolabs, Beverly, MA). The digested DNA was resolved on an ethidium bromide-stained agarose gel (2%). For sequencing, PCR products were cloned using the DynaExpress TA PCR cloning kit (BioDynamics Laboratory Inc, Tokyo, Japan), and the mutations were identified by Sanger sequencing.

### Skeletal staining

Alizarin red and alcian blue staining was performed, as previously described (McLeod, 1980). Samples (postnatal day 1 mice) were fixed in 95% ethanol for 1 week, placed in acetone for 2 d, and then incubated with 0.015% alcian blue 8GS, 0.005% alizarin red S, and 5% acetic acid in 70% ethanol for 3 d. After washing in distilled water, the samples were cleared in 1% KOH for at least 2 d and then in 1% KOH glycerol series until the surrounding tissues turned transparent. The specimens were stored in glycerol until morphological analysis under a `stereomicroscope`.

### Dissection of the anterior and posterior parts of the embryo

Dissection of the anterior and posterior parts of the embryo (referred to as the “head” and “trunk”, respectively) was performed as described previously (Kondrashov et al., 2011; Terranova et al., 2006), with minor modifications. Briefly, the wild-type and *Kdm7a*^−/−^ embryos were staged precisely by counting the somites. Embryos at somite stage 25 (E9.5) and 40 (E10.5) were used for the majority of experiments in the current study. The embryonic head and trunk were divided at the level of otic vesicle, by utilizing micro-surgical scissors. The embryonic head and trunk were then transferred directly to QIAzol® lysis reagent (Qiagen, Hilden, Germany) and were stored at −80°C for RNA isolation.

### mRNA isolation

Total RNA was isolated from the embryonic head and trunk by using a miRNeasy micro kit (Qiagen) with the DNase digestion step, according to the manufacturer’s instructions.

### qPCR for mRNA quantification

The isolated RNA (500 ng) was reverse-transcribed to cDNA by using PrimeScript RT master mix (Takara, Shiga, Japan). PCR was performed using a CFX96 unit (Bio-Rad, Hercules, CA) with SYBR® Premix EX Taq™ II (Takara). The relative expression levels were calculated using *β*-*actin* mRNA as a reference. The primers used for these analyses are listed in Supplementary Table 3.

### ChIP-qPCR

The embryonic head and trunk were collected as described in the section *Dissection of the anterior and posterior parts of the embryo*. To prepare single cell suspension, the tissues were placed in 1 ml of phosphate-buffered saline, pipetted and passed through a 35-μm cell strainer (Corning Japan, Tokyo, Japan). The cells were fixed for 10 min in a 1% formaldehyde solution at room temperature and then neutralized for 5 min in a 0.125 M glycine solution. Pooled tissue samples from two embryos were used in ChIP analysis. ChIP was performed as previously described (Kanki et al., 2011; Kanki et al., 2017). Briefly, fixed cells were re-suspended in 2 ml of sodium dodecyl sulfate lysis buffer, containing 10 mM Tris-HCl, pH 8.0 (Thermo Fisher Scientific), 150 mM NaCl (Thermo Fisher Scientific), 1% sodium dodecyl sulfate (Sigma-Aldrich, St. Louis, MO), 1 mM EDTA, pH 8.0 (Thermo Fisher Scientific), and cOmplete™ EDTA-free protease inhibitor cocktail (Sigma-Aldrich). The samples were then fragmented in a Picoruptor (40 cycles, 30 s on/30 s off; Diagenode, Liege Science Park, Belgium). The sonicated solution was diluted with ChIP dilution buffer [20 mM Tris-HCl, pH 8.0, 150 mM NaCl, 1 mM EDTA, and 1% Triton X-100 (Sigma-Aldrich)] up to 10.3 ml; 10 ml were used for immunoprecipitation (10 ml) and the remaining 300 μl were saved as a non-immunoprecipitated chromatin (the input sample). Specific antibodies against H3K4me3 and H3K9me2 (MAB Institute, Inc. Nagano, Japan) were bound to magnetic Dynabeads M-280 (Thermo Fisher Scientific) and mixed with the diluted, sonicated solution for immunoprecipitation. The prepared DNA was quantified using a NanoDrop 2000 spectrophotometer (Thermo Fisher Scientific), and more than 10 ng of DNA were processed for qPCR. The quantification primers are listed in Supplementary Table 4. PCR was performed using a CFX96 PCR and SYBR® Premix EX Taq™ II. Fold enrichment was determined as the percentage of the input.

### ChIP-Seq library preparation

ChIP-Seq library was prepared using DNA sonicated to an average size of 0.5◻kb. ChIP samples were processed for library preparation using a KAPA Hyper Prep kit (Kapa Biosystems Inc., Wilmington, MA), according to the manufacturer’s instructions. Deep sequencing was performed using a HiSeq 2500 sequencer (Illumina Inc., San Diego, CA) as single-end 36-b reads.

### RNA-Seq library preparation

Total RNA from the embryos was isolated as described above in the section *mRNA isolation*. The RNA integrity score was calculated using the RNA 6000 Nano reagent (Agilent Technologies) and a 2100 Bioanalyzer (Agilent Technologies). RNA integrity value (RIN) score of all samples used for the preparation of RNA-Seq libraries was above 9. RNA-Seq libraries were prepared with a TruSeq RNA Library Prep Kit (Illumina). The libraries were sequenced using a HiSeq 2500 sequencer (Illumina) as paired-end 150-b reads.

### Bioinformatics

#### RNA-Seq data analysis

The quality of FASTQ files was checked by using FastQC (http://www.bioinformatics.babraham.ac.uk/projects/fastqc) version 0.11.8, and trimmed using Trimmomatic PE version 0.38 (Bolger et al., 2014) with “ILLUMINACLIP:adaptor_sequence.fa:2:30:7:1:true LEADING:3 TRAILING:3 SLIDINGWINDOW:4:15 CROP:120 MINLEN:36” parameters. The trimmed FASTQ files were aligned to the mouse reference genome mm10 using Hisat2 version 2.1.0 (Kim et al., 2015) with a “--dta” option. SAM files were sorted and converted into BAM files using Samtools version 1.9 (Li et al., 2009). Gene expression was quantified in transcripts per lilobase million (TPM) using StringTie version 1.3.4d (Pertea et al., 2015) with an “-e” parameter; the GTF file was downloaded from GENCODE release M20 (https://www.gencodegenes.org/mouse/release_M20.html) and input with a “-G” option. To visualize the sequencing tracks, BIGWIG files were generated from BAM files using deepTools version 3.2.0 (Ramirez et al., 2016), bamCoverage command with “-of bigwig -bs 1 --exactScaling --normalizeUsing CPM” parameters, and displayed in Integrative Genomics Viewer (Robinson et al., 2011). Read count table was produced using featureCounts version 1.6.3 with “-t exon -g gene_id --extraAttributes gene_name -M -s 0 -p -P -d 0 -D 500 -a gencode.vM20.annotation.gtf” parameters. Differential expression was determined using DESeq2 (Love et al., 2014) by testing wild-type versus *Kdm7a*^−/−^ embryos at E9.5 or E10.5. The values obtained from DESeq2 were used to generate a heat map and volcano plots. Differentially expressed genes were defined based on two criteria: (1) padj<0.05 and (2) TPM>1 in either or both wild-type or KO samples, and used for GO analysis in DAVID (Huang da et al., 2009) and IPA (QIAGEN, https://www.qiagenbioinformatics.com/products/ingenuity-pathway-analysis).

#### ChIP-Seq data analysis

The quality of FASTQ files was by using FastQC (http://www.bioinformatics.babraham.ac.uk/projects/fastqc) version 0.11.8, and trimmed using Trimmomatic SE version 0.38 (Bolger et al., 2014) with “ILLUMINACLIP:adaptor_sequence.fa:2:30:7 LEADING:3 TRAILING:3 SLIDINGWINDOW:4:15 MINLEN:36” parameters. The trimmed FASTQ files were aligned to the mouse reference genome mm10 using Bowtie2 version 2.3.4.3 (Langmead and Salzberg, 2012) with the default parameters. SAM files were sorted and converted into BAM files using Samtools version 1.9 (Li et al., 2009). To visualize the sequencing tracks, BIGWIG files were generated from BAM files using deepTools version 3.2.0 (Ramirez et al., 2016) bamCoverage command with “-of bigwig -bs 1 --exactScaling --normalizeUsing CPM -e 300” (H3K4me3, and input control), or “-of bigwig -bs 1 --exactScaling --normalizeUsing CPM -e 500” (H3K27me2) parameters, and displayed in Integrative Genomics Viewer (Robinson et al., 2011). For calculating log2 fold-change of the ChIP-Seq signal, sequence reads for the *Hox* family genes were counted using the featureCounts version 1.6.3 with “-t gene -g gene_id --readExtension3 300 --extraAttributes gene_name -M -O -s 0 -a gencode.vM20.annotation.gtf” (H3K4me3) or “-t gene -g gene_id --readExtension3 500 --extraAttributes gene_name -M -O -s 0 -a gencode.vM20.annotation.gtf” (H3K9me2) parameters. Read count for each *Hox* gene was normalized to the total number of mapped reads (per million mapped reads) and gene cluster-specific background noise, which is the total number of input reads for *Hoxa*, *Hoxb*, *Hoxc*, or *Hoxd* genes.

### QUANTIFICATION AND STATISTICAL ANALYSIS

Statistical differences were analyzed by using the Student’s *t*-test. In all tests, differences at *P*-values of <0.05 were considered to be statistically significant.

## DATA AND CODE AVAILABILITY

Sequence data can be accessed through the Gene Expression Omnibus (GEO) under the NCBI accession number GSE133188.

## SUPPLEMENTAL ITEMS

Table S1. Summary of RNA-Seq analysis, related to Figure 2

Table S2. List of primers and oligonucleotides used for construct generation

Table S3. Primers used for mRNA quantification

Table S4. Primers used for ChIP followed by qPCR

