## Supplemental Table 2 for "Lysine demethylase 7a regulates the anterior-posterior development in mouse by modulating the transcription of *Hox* gene cluster"

Supplementary Table2: List of primers and oligonucleotides used for construct generation.

|  | Sequence |
| --- | --- |
| Kdm7a_51+_for_pCAG-EGxxFP | AGCTACATAATGAAACCATGTCTCC |
| Kdm7a_564-_for_pCAG-EGxxFP | GTACCAAGGCAGAAAATGTTTAAGA |
| sgRNA_867F_for_pX330 | CACCGTATAGCCAGAAAACTTTCA |
| sgRNA_867R_for_pX330 | AAACTGAAAGTTTTCTGGCTATAC |
| sgRNA_868F_for_pX330 | CACCGATAGCCAGAAAACTTTCAT |
| sgRNA_868R_for_pX330 | AAACATGAAAGTTTTCTGGCTATC |
| sgRNA_868F_for_DR274 | TAGGTATAGCCAGAAAACTTTCAT |
| sgRNA_868R_for_DR274 | AAACATGAAAGTTTTCTGGCTATC |
