## Supplemental Table 3 for "Lysine demethylase 7a regulates the anterior-posterior development in mouse by modulating the transcription of *Hox* gene cluster"

Supplementary Table 3: Primers used for mRNA quantification

| Target gene | Strand | Sequence (5′ end) |
| --- | --- | --- |
| Hoxa1 | Forward  Reverse | CTCCCAAAACAGGGAAAGTTGGA  TTGAAGTGGAACTCCTTCTCCAG |
| Hoxb1 | Forward  Reverse | CCCACCTAAGACAGCGAAGG  TGGTGAAGTTTGTGCGGAGA |
| Hoxd1 | Forward  Reverse | CCCCCAAGAAAAGCAAACTGTC  GGTTCTGGAACCAGATTTTGACC |
| Hoxa2 | Forward  Reverse | CCACAAAGAATCCCTGGAAATAGC  TCACTTGTCTCTCGGTCAAATCC |
| Hoxb2 | Forward  Reverse | GCGAAATTGCTCCATTGCATAAAC  ACCCAATCTCCCTCTCAAATTCAA |
| Hoxa3 | Forward  Reverse | TTAGGTCCAGAAGTGTCCAAACC  CAGTGTCCAGGCACTCTTAACAT |
| Hoxb3 | Forward  Reverse | AAAAAGTGTTAGCCGTCTCTCCG  CGAGAAATCTCCCCTCCTCTGA |
| Hoxd3 | Forward  Reverse | TTCCACTTCAACCGCTATCTGTG  GGAGAATGCAGGATGCCCTTAG |
| Hoxa4 | Forward  Reverse | GAGCGCCGTCAACTCCAGTTAT  AGTGGAATTCCTTCTCCAGTTCC |
| Hoxb4 | Forward  Reverse | GAGCACGGTAAACCCCAATTACG  GAAACTCCTTCTCCAACTCCAGG |
| Hoxc4 | Forward  Reverse | AGCACGGTGAACCCCAATTA  CGATCTCGATCCTTCTCCTTCG |
| Hoxd4 | Forward  Reverse | TGTAGCGAGCAGCAATACTTAGA  AGTGAATTCTCCTACAAGCCTGG |
| Hoxa5 | Forward  Reverse | AGTGAATTCTCCTACAAGCCTGG  AGTGGAATTCTTTCTCCAGCTCC |
| Hoxb5 | Forward  Reverse | CTTCACATCAGCCACGATATGAC  GGAACCAGATTTTGATCTGACGC |
| Hoxc5 | Forward  Reverse | ACATGAGCCACGAGACGGAT  ATTCTTTCTCGAGTTCCAGGGTC |
| Hoxa6 | Forward  Reverse | GGGACTACCTGCACTTTTCTCC  ATACACGGCACCCGCACAG |
| Hoxb6 | Forward  Reverse | TGAATTCGTGCAACAGTTCCTCT  CCGGTTCTGAAACCAAATCTTGA |
| Hoxc6 | Forward  Reverse | GAATGAATTCGCACAGTGGGGTC  GAGTTAGGTAGCGGTTGAAGTGA |
| Hoxa7 | Forward  Reverse | AGTTCAGGACCCGACAGGAAG  TGGAATTCCTTCTCCAGTTCCAG |
| Hoxb7 | Forward  Reverse | TCAAGGAATCTCGTAAAACCGAC  TGAACTCATAATTTGGCCGGATG |
| Hoxb8 | Forward  Reverse | GGACCTTTTAAAACTCGGTGCAA  TCTTTCTAAATGTCAGGGTCGCT |
| Hoxc8 | Forward  Reverse | GGATGAGACCCCACGCTCC  CTTGTCTTTCTGTCAGTCCCAGG |
| Hoxd8 | Forward  Reverse | CCGCGAAGTTTTACGGATACGAT  GGAGCTGCTTGTGGTCTCATC |
| Hoxa9 | Forward  Reverse | GCATTAAACCTGAACCGCTCTC  CGGGTTATTGGGATCGATGGG |
| Hoxb9 | Forward  Reverse | AAAGAGAGGCCGGATCAAACCAA  GGTCCCTGGTGAGGTACATATTG |
| Hoxc9 | Forward  Reverse | AAAAGATCAGAGACTGCAGGAGC  GATGAAAATGCCAGTCCCAGAAG |
| Hoxd9 | Forward  Reverse | CAACTTGACCCAAACAACCCTG  CTCTAGCGTCTGGTATTTGGTGT |
| Hoxa10 | Forward  Reverse | GAGTCCTAGACTCCACGCCA  CCTTTGGAACTGCCCAGGGA |
| Hoxc10 | Forward  Reverse | CGGATAACGAAGCGAAAGAGGAG  AATGGTCTTGCTAATCTCCAGGC |
| Hoxd10 | Forward  Reverse | CAGGAGAAGGAAAGCAAAGAGGA  GGTGAGGTTAACGCTCTTACTGA |
| Hoxa11 | Forward  Reverse | ATATCATCCCACCACTGATCTGC  CACAGCCTCTGGAGTTTTCAATG |
| Hoxc11 | Forward  Reverse | ATGTTTAACTCGGTCAACCTGGG  TAAGTGCAACTGGGCAGATAGAG |
| Hoxd11 | Forward  Reverse | GGCGAGATCTGTAGGAAGTTAGG  CCCAAAGGTACATTTCCCAGAGT |
| Hoxc12 | Forward  Reverse | CCTACTCAACGAGGGCAATAAGA  TGATGAACTCGTTGACCAGAAAC |
| Hoxd12 | Forward  Reverse | CCAACCTTTAGCAAGATGCACAA  ACATAAACGGCAACTGTTAGCAC |
| Hoxa13 | Forward  Reverse | GAAAGAACTCGAACGGGAATACG  CTCCTGTTCTGGAACCAGATTGT |
| Hoxc13 | Forward  Reverse | CAGTCAGGTGTACTGCTCCAAG  TCTTTGGTGATGAATTTGCTGGC |
| Hoxd13 | Forward  Reverse | TCCTTTCCAGGAGATGTGGCT  TCTCTCCGAAAGGTTCGTGG |
| Kdm7a | Forward  Reverse | AGCAATAGAGGAGGAAAATGGCA  CAAGGTTAGAAGGAGTTCGGACA |
