## Supplemental Table 4 for "Lysine demethylase 7a regulates the anterior-posterior development in mouse by modulating the transcription of *Hox* gene cluster"

|  | Strand | Sequence (5′ end) |
| --- | --- | --- |
| Actb | Forward  Reverse | GCCGTATTAGGTCCATCTTGAGA  CAAACCGGTTTGGACAAAGACC |
| Hoxa3 | Forward  Reverse | TCGTGCTGCTAAATATTGCTGAC  GCGCAAATCCATCTTACTCTCAA |
| Hoxa13 | Forward  Reverse | TCCCTAAAACATGCCAGGACATC  AGTCAGGTAAATTCTCCAGTGGC |

Supplementary Table 4: Primers used for ChIP followed by real-time PCR
